# The effect of assortative mixing on stability of low helminth transmission levels and on the impact of mass drug administration: model explorations for onchocerciasis

**DOI:** 10.1101/354084

**Authors:** Anneke S. de Vos, Wilma A. Stolk, Sake J. de Vlas, Luc E. Coffeng

## Abstract

**Background:** Stable low pre-control prevalences of helminth infection are not uncommon in field settings, yet it is poorly understood how such low levels can be sustained, thereby challenging efforts to model them. Disentangling possible facilitating mechanisms is important, since these may differently affect intervention impact. Here we explore the role of assortative (i.e. non-homogenous) mixing and exposure heterogeneity in helminth transmission, using onchocerciasis as an example.

**Methodology/Principal Findings:** We extended the established individual-based model ONCHOSIM to allow for assortative mixing, assuming that individuals who are relatively more exposed to fly bites are more connected to each other than other individuals in the population as a result of differential exposure to a sub-population of blackflies. We used the model to investigate how transmission stability, equilibrium microfilariae (mf) prevalence and intensity, and impact of mass drug administration depend on the assumed degree of assortative mixing and exposure heterogeneity, for a typical rural population of about 400 individuals. The model clearly demonstrated that with homogeneous mixing and moderate levels of exposure heterogeneity, onchocerciasis could not be sustained below 35% mf prevalence. In contrast, assortative mixing stabilised onchocerciasis prevalence at levels as low as 8% mf prevalence. Increasing levels of assortative mixing significantly reduced the probability of interrupting transmission, given the same duration and coverage of mass drug administration.

**Conclusions/Significance:** Assortative mixing patterns are an important factor to explain stable low prevalence situations and are highly relevant for prospects of elimination. Their effect on the pre-control distribution of mf intensities in human populations is only detectable in settings with mf prevalences <30%, where high skin mf density in mf-positive people may be an indication of assortative mixing. Local spatial variation in larval infection intensity in the blackfly intermediate host may also be an indicator of assortative mixing.

**Author summary:** Most mathematical models for parasitic worm infections predict that at low prevalences transmission will fade out spontaneously because of the low mating probability of male and female worms. However, sustained low prevalence situations do exist in reality. Low prevalence areas have become of particular interest now that several worm infections are being targeted for elimination and the question arises whether transmission in such areas is driven locally and should be targeted with interventions. We hypothesise that an explanation for the existence of low prevalence areas is assortative mixing, which is the preferential mixing of high-risk groups among themselves and which has been shown to play an important role in transmission of other infectious diseases. For onchocerciasis, assortative mixing would mean that transmission is sustained by a sub-group of people and a connected sub-population of the blackfly intermediate host that mix preferentially with each other. Using a mathematical model, we study how assortative mixing allows for sustained low prevalences and show that it decreases the probability of interrupting transmission by means of mass drug administration. We further identify data sources that may be used to quantify the degree of assortative mixing in field settings.

## Introduction

Onchocerciasis prevalence varies widely between geographical locations, with nodule and microfiladermia (mf) prevalence levels in adults ranging from just above 0% to over 80% [1,2]. Onchocerciasis control programmes historically aimed for morbidity control and focussed interventions on so-called meso and hyperendemic areas, i.e. areas with mf prevalence levels above 40%. Many hypoendemic areas (mf prevalence <40%) were left untreated [3]. Now the target has shifted to elimination the question has arisen whether such hypoendemic areas can maintain themselves and may act as a source of infection for areas that have achieved elimination. If so, hypoendemic areas should be covered by elimination campaigns. Answering these questions is not straightforward, as the transmission dynamics in hypoendemic settings are not fully understood. This also applies to other helminthic diseases that are currently the subject of large-scale control and elimination programmes, such as lymphatic filariasis (LF), schistosomiasis and soil-transmitted helminthiasis.

Mathematical models can be useful tools to understand how various processes can help to stabilize helminth transmission in low endemic areas. Population dynamics of helminth infections are unique given the need for male and female worms to be present in the same host for reproduction, leading to a so-called breakpoint prevalence below which transmission cannot maintain itself [4,5]. Most models for helminth transmission explain sustained low pre-control prevalences by assuming high degrees of exposure heterogeneity among human hosts [6–10], meaning that some people are heavily exposed while the majority experience much lower exposure levels. The resulting concentration of worms in few heavily exposed individuals allows female and male worms to mate, even if overall worm numbers in the host population are low. In addition, existing models for helminth transmission typically assume homogeneous mixing. This assumption implies that every person can infect any other person in the community with probability directly proportional to the product of one person’s contribution and another person’s exposure to transmission, as if all transmission takes place in a singular point in space. However, in reality mixing patterns in helminth transmission are assortative (i.e. non-homogeneous) as sub-groups of human hosts mix preferentially and transmit infection amongst themselves because they spend different amounts of time in different shared locations such as e.g. schools, water collection sites, and/or household locations. In summary, assortative mixing in helminth transmission implies the existence of multiple vector or environmental reservoirs and differential exposure of individuals to such reservoirs with a sub-group of high-risk individuals concentrating around at least one of those reservoirs, which is very well conceivable.

Here, we consider for the first time to which extent assortative mixing may play a role in sustaining low levels of helminth transmission. Assortative mixing has been shown to play an important role in the transmission of many infections [11–15]. Especially for sexually transmitted or drug-use related infections, individuals often infect those of similar risk level to their own, as they meet at specific venues or parties [13,14]. In onchocerciasis transmission, which we consider here, there may be specific sub-groups of humans spending relatively much time where fly densities are highest; for example, fisherman will be often near the water where fly breeding sites are found [1]. It is very well conceivable that these high-risk individuals would not only be bitten more often (as assumed by current models), but also more often by flies that previously bit another (or the same) high-risk individual. Under this assumption, the probability of infections spilling over from the highly exposed fishermen to the rest of the community is relatively lower, which means that in very low endemic situations transmission events are not “wasted” on transmission from fishermen to the rest to the population, but more efficiently used to sustain a high concentration of worms in the fishermen, sustaining transmission at relatively low prevalence.

In this paper, we explore how adding assortative mixing to the individual-based model ONCHOSIM impacts onchocerciasis equilibrium prevalence levels and can explain stable low prevalence levels. Furthermore, we show how the (combination of) mechanisms for sustaining low prevalence will be relevant for the impact of control measures, especially when pushing for elimination. Having shown its potential importance, we consider what field data might enable us to identify and quantify assortative mixing in field situations. The findings of our study are also of relevance for other helminth infections that require mating of male and female worms.

## Methods

We use the model ONCHOSIM, an established individual-based model for transmission and control of onchocerciasis [16–21]. ONCHOSIM simulates the individual life histories of humans and the male and female worms living within them. Patent female worms produce microfilariae (mf) as long as there is at least one patent male worm present in the same host. Flies biting on hosts take up mf, but their uptake capacity is limited resulting in diminishing returns with increasing mf levels in hosts (i.e. negative density dependence). Individual human exposure to fly bites is assumed to vary with age and sex, and to vary randomly between individuals as a consequence of other factors (e.g. attractiveness, occupation), leading to a highly overdispersed worm population within the human population. The model further simulates the impact of treatment with ivermectin in context of a mass drug administration, accounting for variation in participation by age and sex and presence of potential systematic non-participation by a subset of individuals. Ivermectin is assumed to kill all microfilariae in treated individuals and to permanently reduce the reproductive capacity of adult female worms by 35%, allowing for cumulative effects of repeated treatments. In addition, after treatment female worms temporarily stop producing mf but gradually recover to their new maximum reproductive capacity in a period of 11 months on average. The model provides output in terms of simulated skin snip surveys (two snips per person), assuming that all individuals in the population are sampled. More technical details and quantification of the “default” model (i.e. with homogeneous mixing) can be found elsewhere [20]. To investigate the effect of assortative mixing on pre-control equilibrium prevalence and intervention impact, the default model was reprogrammed in R and extended as follows.

In the default model, the fly vector population is represented as a single fly population that transmits infectious material (larvae) from human to human. To simulate assortative mixing we have divided this fly population into two sub-populations, which we name fly population *L* and *H* that are relatively more connected with low and high risk groups of the human population, respectively. As in the default model, an individual’s exposure to fly bites is determined by his or her age, gender, and a lifelong relative exposure factor *γ*_*i*_ that represents variation due to random factors such as occupation and attractiveness for flies; *γ*_*i*_ is drawn from a gamma distribution with shape and rate equal to *k* (i.e. mean = 1.0). S1 Figure illustrates the assumed distribution of individual relative exposure under the default assumption of *k* = 3.5 (used in previous ONCHOSIM modelling studies) and an alternative scenario with a higher level of exposure heterogeneity of *k* = 1.0, which we consider to be still realistic and relevant for low endemic situations [19]. For each human *i* we define that his or her vector contacts are divided between the two fly sub-populations as a function of *γ*_*i*_ such that those who are bitten less often are bitten mostly by flies from population *L*, and vice versa those with high exposure to fly bites are bitten most often by flies from population *H*. This leads to assortative mixing, i.e. greater connectedness of individuals with similar risk levels.

We define the fraction of an individual’s total fly contacts that are with fly population *H* (rather than with fly population *L*) as a function of an individual’s relative exposure in terms of his or her percentile *r*(*γ*_*i*_) relative to the rest of the population: *B*-iCDF(*x* = *r*(*γ*_*i*_) |*α*, *β*). Here *B*-iCDF is the inverse-cumulative beta distribution function (naturally bounded between 0 and 1) with shape parameters *α* and *β* and *r*(.) is the cumulative gamma distribution function with shape and rate equal to *k*, the model parameter for exposure heterogeneity. We further set *α* = (1 − *s*) / *s* and *β* = ((1 − *s*)/*s*) ∙ *S*, where *s* (range 0-1) scales the strength of segregation between the two groups (steepness of the population connection distributional curve in S2 Figure) and *S* is solved numerically such that *B*-iCDF (*x* = *f*_*H*_ | *α*, *β*) = 0.5, where *f*_*H*_ is the parameter for the proportion of the population that is relatively more exposed to fly population *H* (i.e. more than 50% of these individuals’ contacts with flies are with flies from fly population *H*). S2 Figure illustrates the association between individual relative exposure and different fractions of fly contacts with fly population *H* considered in this paper (*f_H_* = 0.5, 0.25 and 0.1).

When *s* = 1 we have two fully separate pairs of human and fly populations. When *s* <1, the association between individual relative exposure and fraction of bites received from fly population H follows an s-curve (S2 Figure), with higher steepness in the middle for higher values of *s*. When *s* = 0, the fraction of fly contacts that an individual has with flies from fly population *H* is the same (i.e. *f_H_*) for all individuals, resulting in homogenous mixing. For illustrative purposes, we only consider relatively strong assortative mixing (*s* = 0.8). For the homogenous mixing scenario, we compare medium (*k* = 3.5) with high (*k* = 1) heterogeneity in individual exposure to fly bites. Note that the fraction of all fly bites that are from fly population *H* will be substantially larger than the fraction of humans *f_H_* connected mostly to fly population *H*: when *k* = 3.5*, s* = 0.8, and *f_H_* respectively 0.5, 0.25 and 0.1, the fraction of all bites by flies from population *H* is 69%, 44% and 26% (see also S3 Figure).

The model concepts for assortative mixing described above were implemented in a new version of the original model [20] which we programmed in R. We simplified the R version of the model for a limited number of factors that we consider to be of minor relevance to the research question investigated here. First, the model does not distinguish between male and female humans and therefore assumes no difference in exposure to fly bites between the sexes. Second, survival of microfilariae is assumed to be exponential instead of having a fixed duration, which is of limited importance when comparing the impact of MDA (which kills microfilariae) under different assumptions about mixing patterns. Third, we do not consider a fraction of individuals that are permanently excluded from MDA due to pre-existing conditions, nor do we consider non-participation due to e.g. pregnancy (i.e. everybody is eligible for treatment). We do however only allow individuals of age five and above to be treated in MDA, as before. Fourth, all worms and humans are always born at the start of each monthly time step in the model, instead of spread out over the month. Finally, to explore the potential impact of random vs. systematic MDA participation, we included the model concept recently developed by Irvine et al. [9], which is more parsimonious compared to that in ONCHOSIM. With these simplifications, the R version of the ONCHOSIM could very closely reproduce predictions in terms of prevalence and intensity of infection by the original model.

## Results

Figure 1 shows how the mean annual fly biting rate (ABR) determines the dynamic equilibrium mf prevalence level at which onchocerciasis transmission is sustained in the absence of interventions. At a moderate level of heterogeneity in individual exposure to fly bites (scenario “*k* = 3.5 (one fly population)”, i.e. the default assumption in previous ONCHOSIM modelling studies), we see a very steep decline in equilibrium skin microfilarial (mf) prevalence with decreased ABR, especially at ABR below 12,000. At around ABR = 10,000 we find a boundary in transmission stability (defined as <50% probability of extinction during 200 years of simulation time), which is due to a relative low worm mating probability at lower prevalence combined with the assumed transmission conditions.

**Figure 1.**
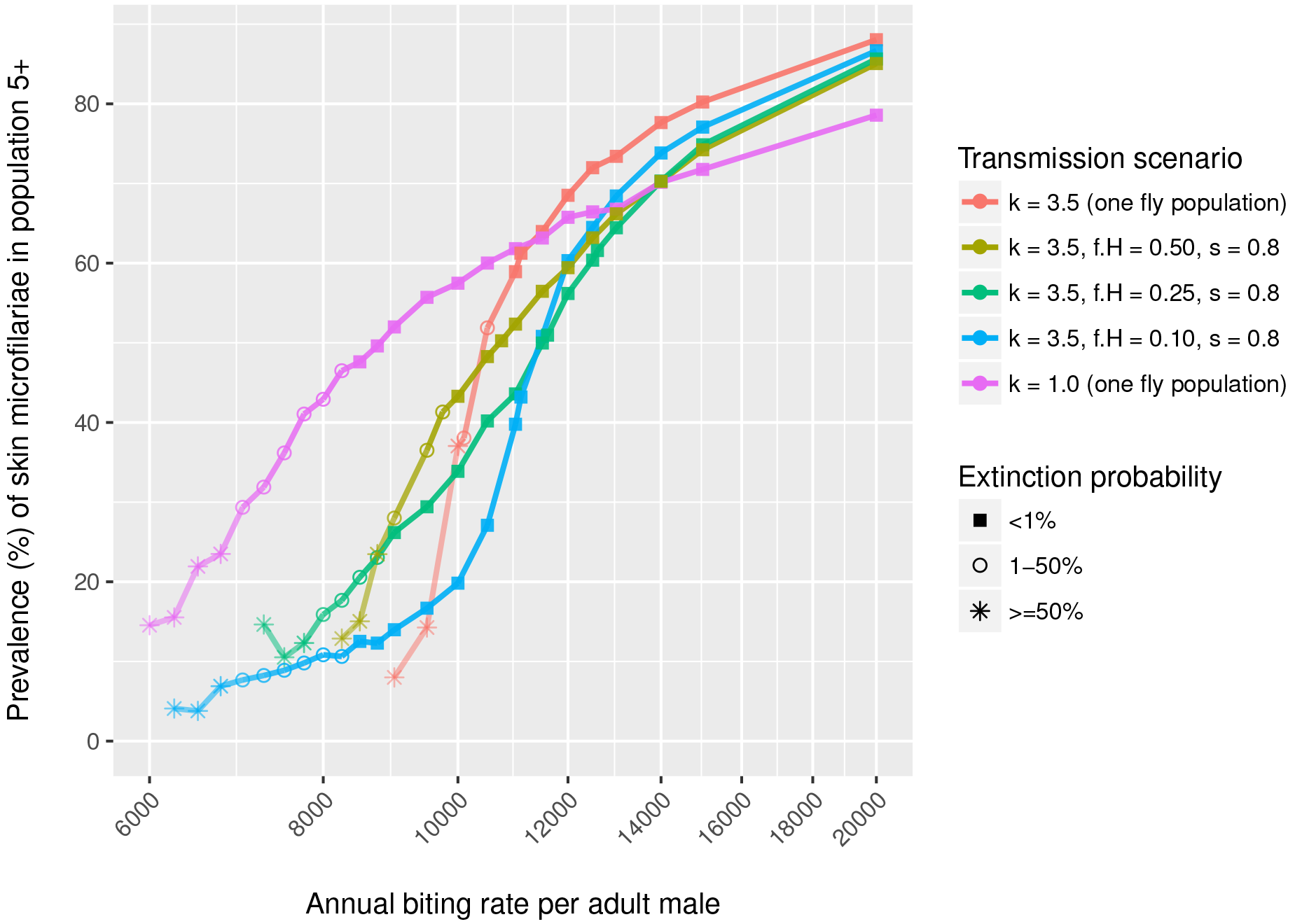
Model-predicted association between annual biting rate, prevalence of skin microfilariae, and stability of transmission. Bullets represent the average skin mf prevalence over 150 repeated simulations, with the shape of the bullet indicating the extinction probability (here defined as the proportion of repeated simulations in which transmission spontaneously faded out within 200 years). The red and purple lines (with k = 3.5 or 1.0 and one fly population) represent transmission scenarios with homogeneous mixing; the other coloured lines represent transmission scenarios with assortative mixing, assuming presence of two fly populations where some proportion *f_H_* of the human population with relative high exposure to flies has most of its contact with the fly population *H*. Parameter *s* represents the level of segregation of the two fly populations, e.g. s = 0 represents homogeneous mixing (presence of two populations but all humans have equal opportunity to be exposed to both) and s = 1 represents two completely segregated fly populations for which the biting affects two completely segregated human populations. See methods section for details.

With greater heterogeneity in individual exposure to fly bites (scenario “*k* = 1.0 (one fly population)”), at a high ABR of 20,000 the achieved mf prevalence decreases from about 88% to 79% (compared to “*k* = 3.5 (one fly population)”). Stronger heterogeneity implies that there is more variation in biting rates experienced by people, resulting in a larger proportion of people with very high number of bites, but also a larger proportion of people experiencing very low number of bites. The latter group has a relatively low risk of infection, which limits the maximum achievable prevalence in the simulation. However, in this more heterogeneous setting the prevalence declines far less steeply with decreasing ABR; that is, transmission remains efficient since those bitten often both carry high worm burdens and they transmit to more flies. As this concentration of worms within fewer individuals allows for continued mating, transmission is now sustained (i.e. probability of extinction <50%) down to mf prevalence of 30%, at an ABR as low as about 7000.

Assortative mixing has less of a dampening impact on prevalence at high biting rates, compared to increasing heterogeneity (i.e. lower values of *k*). Further, it somewhat lowers the threshold ABR below which extinction occurs, but not as much as lower values of *k*. However, it does allow for sustained transmission at much lower biting rates, especially if there is a relatively small higher risk sub-group, whose members are connected through a shared population of vectors. When the high-risk group constitutes 50%, 25% or 10% of the general human population, the model can maintain stable mf prevalences as low as 28%, 16% or even 8%, respectively.

The predicted effect of mass drug administration (MDA) strongly depends on the assumed exposure heterogeneity as well as the mixing pattern within a population (Figure 2). The probability of elimination decreases with higher levels of exposure heterogeneity (purple vs. red lines) and when transmission is concentrated in a smaller part of the population (blue vs. red lines). In case of recrudescence of infection after stopping MDA, the slope of the rebound over time varies highly between simulations in the scenario with homogeneous mixing and high exposure heterogeneity (purple lines), while this variation is much smaller in case of assortative mixing driven by a small fraction of the human population (blue). Also, the speed of bounce-back is slower in the scenario where transmission is concentrated in a smaller subgroup of the general population (blue). These patterns are also seen for other endemicity levels and patterns in MDA participation (S4 Figure).

**Figure 2.**
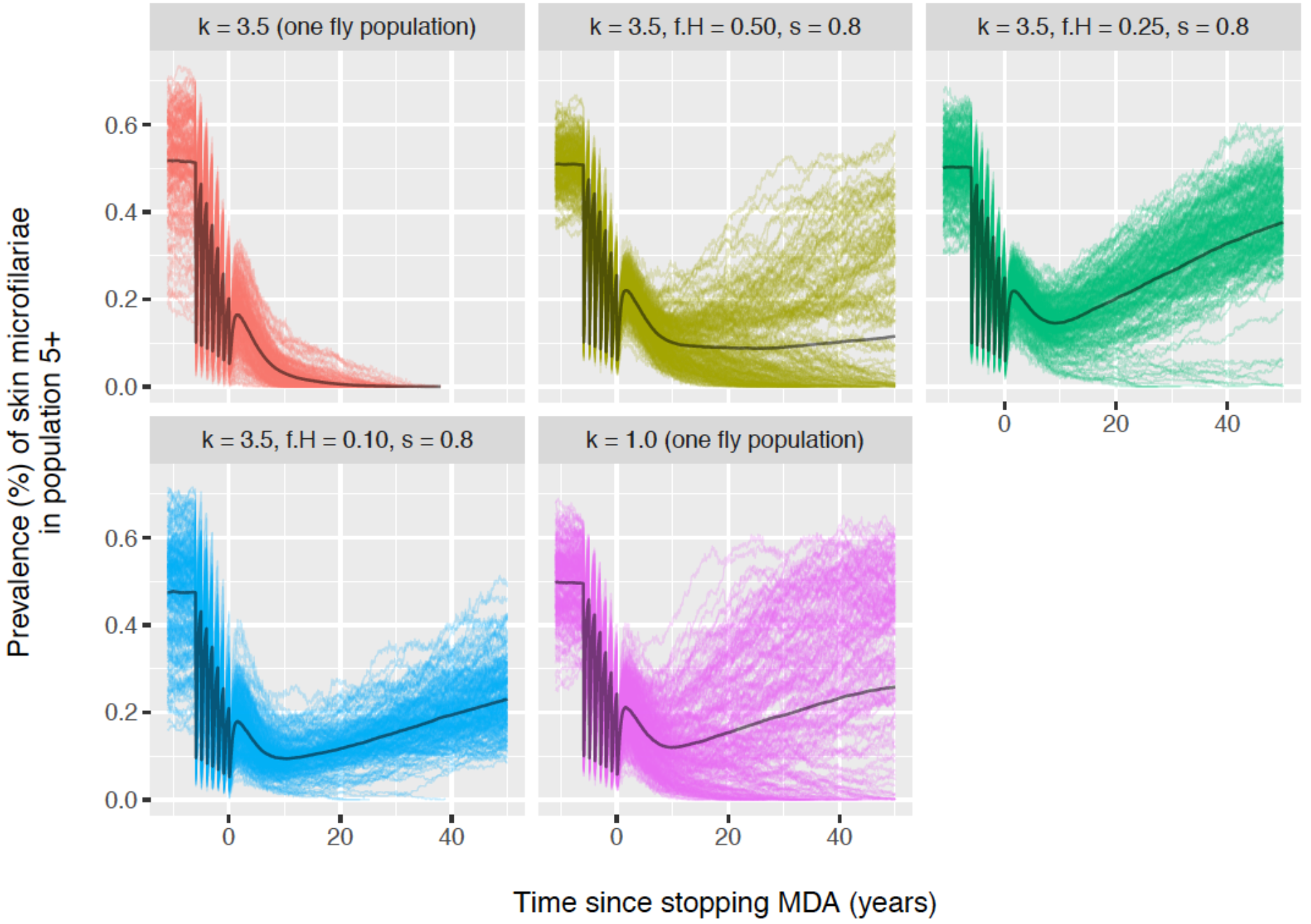
The influence of mixing patterns on trends in prevalence of skin microfilariae during mass drug administration. Lines represent results repeated simulations for a fixed annual biting that was tuned (given exposure heterogeneity *k* and assumed mixing pattern) to result in an average pre-control prevalence of about 50% in the population of age 5 and above. In each simulation, 7 mass drug administration (MDA) rounds are implemented at 65% coverage of the general population. Participation to MDA was assumed to be semi-systematic (some individuals are structurally more likely to participate that others). S4 Figure illustrates similar results for other pre-control endemicity levels and assumed patterns in MDA participation.

Table 1 summarises the outcome of simulated scenarios in terms of the probability of elimination (defined as the proportion of repeated simulations with zero worm prevalence 50 years after stopping MDA), confirming the patterns in Figure 2.

**Table 1.** Impact of mixing patterns on probability of elimination.

| Scenario | Probability of elimination (range 0-1)<sup>a</sup> |  |  |
| --- | --- | --- | --- |
|  | Random MDA participation<sup>b</sup> | Semi-systematic MDA participation<sup>b</sup> | Fully systematic MDA participation<sup>b</sup> |
| *40% pre-control mf prevalence in age 5+ and 5 rounds of annual MDA at 65% coverage* |  |  |  |
| k = 3.5 (one fly population) | 0.98 | 0.97 | 0.85 |
| k = 3.5, f<sub>H</sub> = 0.50, mixt = 0.8 | 0.67 | 0.60 | 0.52 |
| k = 3.5, f<sub>H</sub> = 0.25, mixt = 0.8 | 0.02 | 0.03 | 0.01 |
| k = 3.5, f<sub>H</sub> = 0.10, mixt = 0.8 | 0.01 | 0.00 | 0.00 |
| k = 1.0 (one fly population) | 0.42 | 0.33 | 0.27 |
| *50% pre-control mf prevalence in age 5+ and 7 rounds of annual MDA at 65% coverage* |  |  |  |
| k = 3.5 (one fly population) | 0.99 | 1.00 | 0.93 |
| k = 3.5, f<sub>H</sub> = 0.50, mixt = 0.8 | 0.65 | 0.51 | 0.25 |
| k = 3.5, f<sub>H</sub> = 0.25, mixt = 0.8 | 0.04 | 0.03 | 0.00 |
| k = 3.5, f<sub>H</sub> = 0.10, mixt = 0.8 | 0.02 | 0.02 | 0.01 |
| k = 1.0 (one fly population) | 0.28 | 0.32 | 0.14 |
| *60% pre-control mf prevalence in age 5+ and 11 rounds of annual MDA at 65% coverage* |  |  |  |
| k = 3.5 (one fly population) | 1.00 | 1.00 | 0.98 |
| k = 3.5, f<sub>H</sub> = 0.50, mixt = 0.8 | 0.99 | 0.86 | 0.38 |
| k = 3.5, f<sub>H</sub> = 0.25, mixt = 0.8 | 0.54 | 0.25 | 0.06 |
| k = 3.5, f<sub>H</sub> = 0.10, mixt = 0.8 | 0.26 | 0.21 | 0.08 |
| k = 1.0 (one fly population) | 0.57 | 0.39 | 0.06 |

<sup>a</sup> Elimination is defined as zero mf prevalence 50 years after stopping MDA; probability of elimination is defined as the fraction of 200 repeated simulations that meet aforementioned criterion.
<sup>b</sup> Random MDA participation means that every eligible individual is just as likely to participate; fully systematic participation means that always the same eligible persons participate; semi-systematic participation is a mix of random and fully systematic participation.

Finally we consider what real-world data might help us identify whether low pre-control prevalences are the result of stable low transmission facilitated by either assortative mixing or high exposure heterogeneity, or are the result of a transient decline due to stochastic fade-out. Hypothesising that assortative mixing and high exposure heterogeneity impact the distribution of intensity of infection in different ways, we explore the association between prevalence of skin mf and the arithmetic mean skin mf density in mf positives (Figure 3). At low mf prevalences (<30%) the arithmic mean density of mf in mf-positive individuals is considerably higher in settings with strong assortative mixing (*f_H_* = 0.25 and 0.1) compared to in settings with homogeneous mixing with moderate (*k* = 3.5) to high exposure heterogeneity (*k* = 1.0, which we consider a plausible extreme value). As such, relatively high arithmic mean skin mf loads in mf positive persons in settings with mf prevalence <30% may be an indication of stable transmission facilitated by assortative mixing. For settings with pre-control mf prevalences of 40% to 60%, different mixing conditions and levels of exposure heterogeneity result in very similar associations between arithmic mean skin mf density in mf-positives and the mf prevalence (Figure 3) as well as very similar mf intensity distributions (Figure 4). For settings with mf prevalence >60%, arithmic mean skin mf densities are almost identical for different mixing conditions, but are relatively higher in settings with higher exposure heterogeneity (purple line).

**Figure 3.**
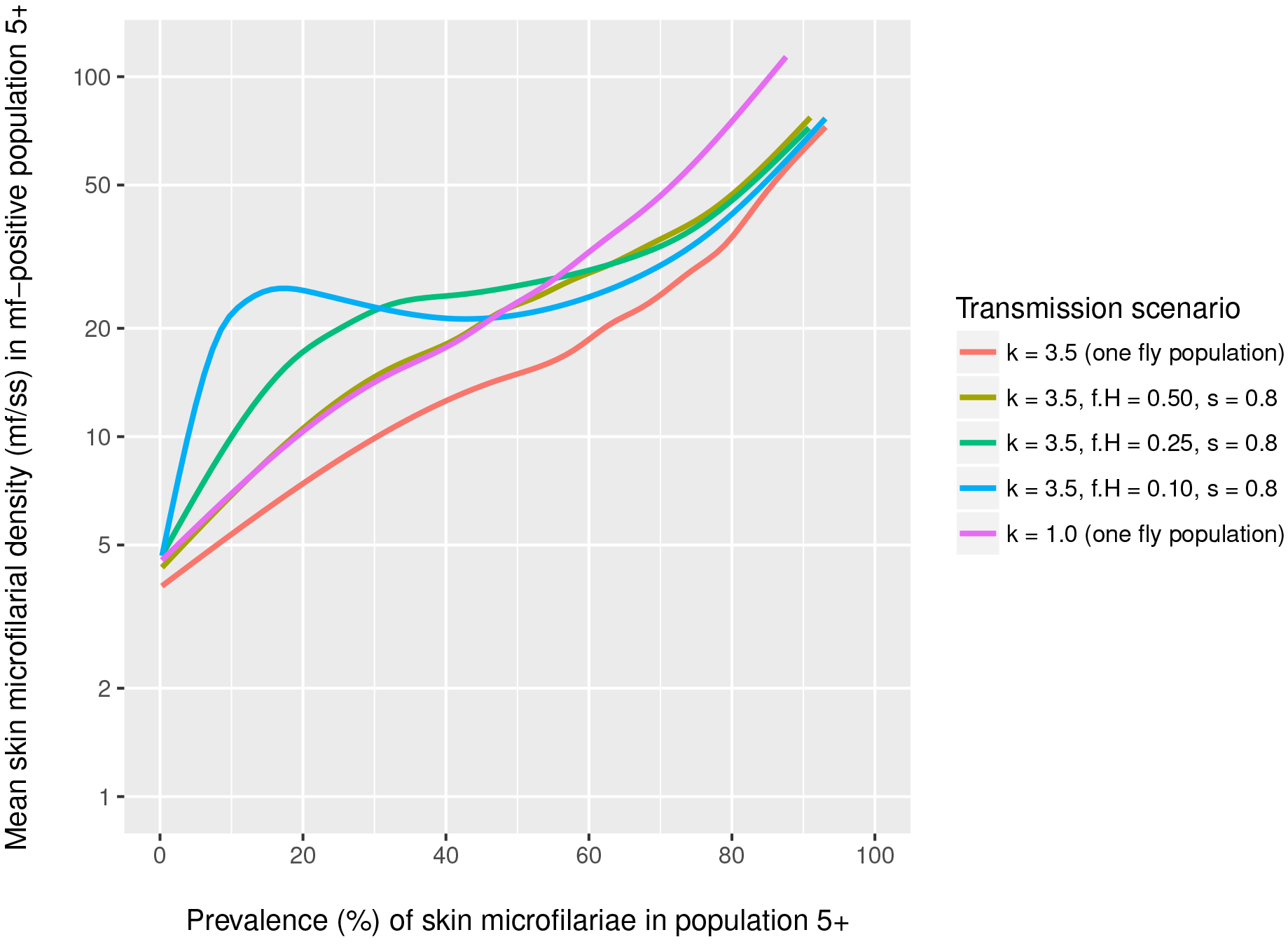
Pre-control arithmic mean density of mf in the skin among mf-positive individuals of age 5+ and above. Lines are based on a generalised additive model with integrated smoothness estimation, fitted to predicted mf prevalences and intensities of repeated simulations for the same range and values of annual biting rate (ABR) used in Figure 1. For each value of ABR 150 repeated simulations were performed. Individual simulation results and the fit of the generalised additive model can be found in S5 Figure. Note that in all scenarios a mean microfilarial density below ~15 mf/ss in the mf-positive population is indicative of stochastic fade-out taking place (i.e. as incidence declines the worm population ages, resulting in lower mf production per female worm).

**Figure 4.**
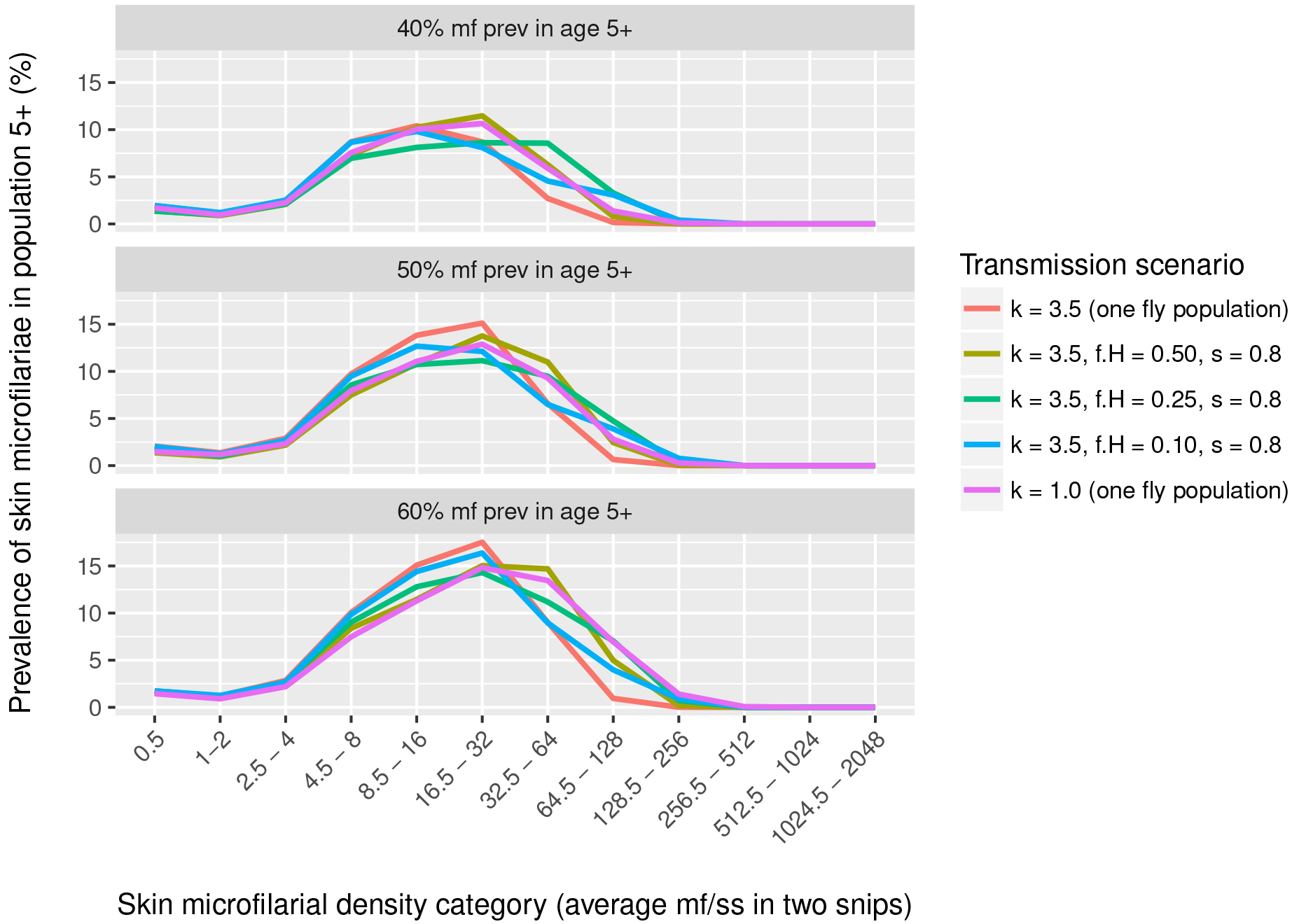
Distribution of skin microfilarial density in mf-positive individuals in different transmission scenarios and endemicity levels. Distributions are based on the average of 150 repeated simulation for each of three fixed values of the annual biting rate that result in an average pre-control mf prevalence of 40%, 50%, and 60% in the population of age 5 and above (three panels).

Another indication for assortative mixing may be found by considering local level fly data, as assortative mixing can only play a role if the mean larval intensity is not equally distributed across fly sub-populations that humans are exposed to. Figure 5 illustrates how the ratio of intensity of infection in the high and low risk fly populations might change with pre-control mf prevalence in humans, assuming perfect measurements from locations with minimal overlap of the two fly populations. A ratio of 1.0 (dashed horizontal black line) represents settings where infection intensity is uniformly distributed across the fly sub-populations (i.e. homogeneous mixing). This ratio increases strongly with lower mf prevalence in humans, with a difference of factor 10 to 50 for settings with mf prevalences under 20%. However, the ratio provides little information about the extent to which transmission is concentrated in a human sub-population (similar curves for different values of *f_H_*).

**Figure 5.**
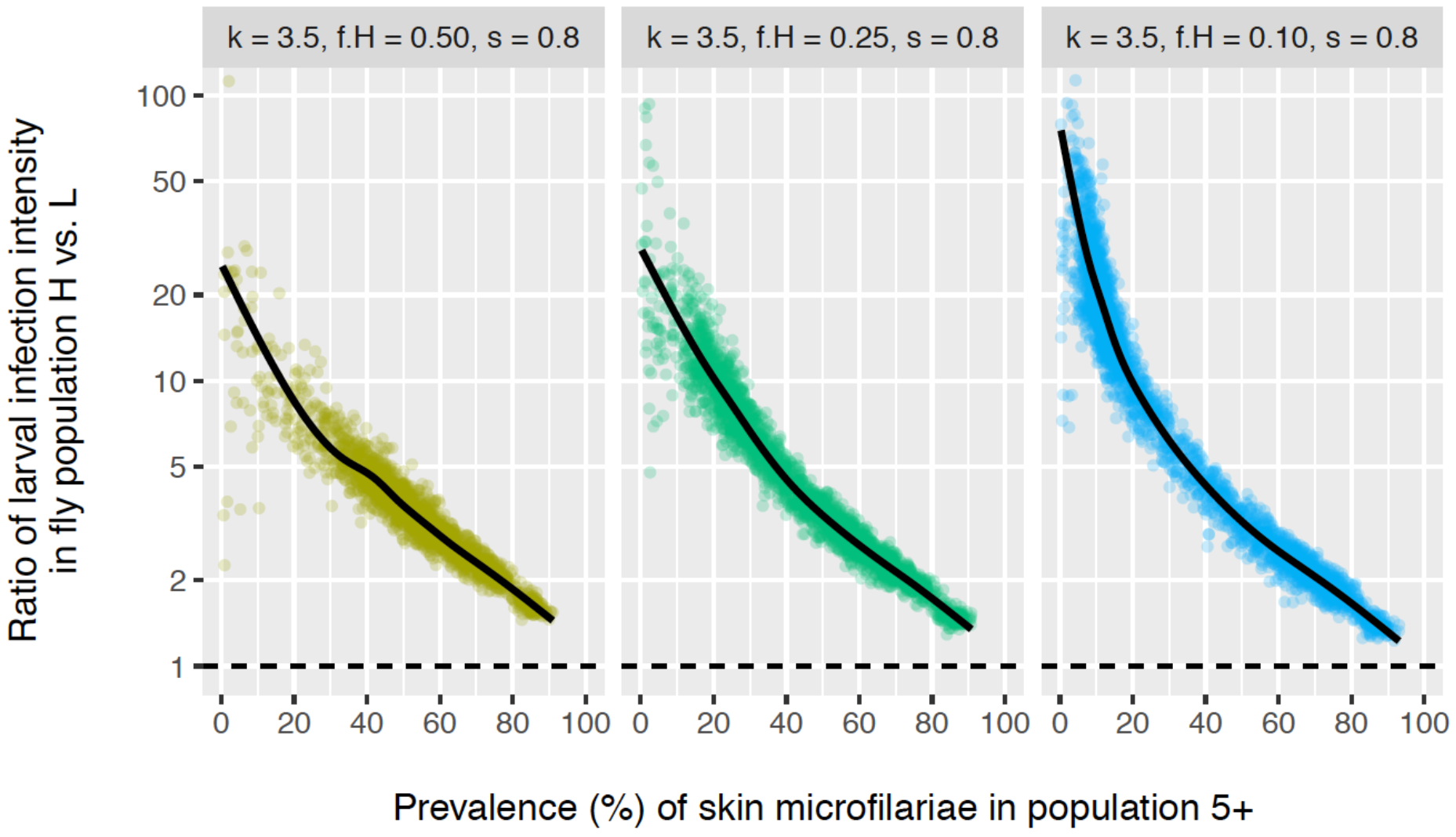
Ratio of larval infection intensity in two spatially separate samples of blackflies around a single community as an indicator of assortative mixing. Each bullets represents the result of a single simulation. Simulations were run using the same range and values of annual biting rate (ABR) as used in Figure 1, and for each value of ABR 150 repeated simulations were performed. For comparison, a ratio close to 1.0 (horizontal dashed black line) would indicate that flies from two spatially separate samples bite humans with a similar distribution of infection levels (i.e. under the assumption of homogeneous mixing). Lines are based on a generalised additive model with integrated smoothness estimation, fitted to individual bullets.

## Discussion

Our study shows that stable low prevalences of onchocerciasis can be explained by both high exposure heterogeneity and assortative mixing. In contrast, if assortative mixing is the main driver of sustained low prevalences, the probability of elimination declines when transmission is sustained by a smaller human sub-population. Also, recrudescence of infection after stopping MDA is slower and less variable in terms of speed when assortative mixing is driven by a smaller human sub-population. Pre-control skin mf density distributions provide little information to distinguish exposure heterogeneity and assortative mixing, or to quantify the degree of assortative mixing. Only in situations with mf prevalence <30%, high arithmic mean skin mf densities (>20 mf/ss) in mf positives may be an indication of assortative mixing. Entomological data may also provide evidence for presence of assortative mixing, but unfortunately not the size of the human sub-population by which it is driven.

Our findings about the role of assortative mixing also apply to the transmission of other human helminth infections. Especially for LF, which is transmitted by mosquitoes and also targeted for elimination, the relatively low mobility of mosquitoes (compared to blackflies) means that people in the same household are likely to be bitten by the same mosquito sub-population near their household [15,22]. In this context, differences between LF vector species mobility and biting behaviour will also be relevant for degree of and patterns in assortative mixing. Similarly, transmission of soil-transmitted helminths and schistosomiasis most likely takes place through multiple reservoirs that are situated near households and/or schools, instead of one central reservoir [23]. Although schistosomiasis and soil-transmitted helminth are not (yet) officially targeted for elimination, there has been increasing interest in the potential of interrupting transmission [10,24–26], which means that also here assortative mixing will become an important factor to consider.

Our study clearly demonstrates that low prevalence of onchocerciasis could be sustained by assortative mixing. Another suggested mechanism to explain low prevalences is that infection spills over from nearby higher endemic areas through movement of infected humans and/or flies [27]. This is undoubtedly true for many of such settings, and can in fact be considered a form of assortative mixing at a wider geographical scale, as it simply constitutes flow of infections between two or more populations with each their own local transmission conditions. As such, we expect that the impact of migration is qualitatively similar to the impact of assortative mixing that we predict here. Another logical alternative explanation of (seemingly stable) low endemic levels is that these are the result of high transmission in the past that has stopped due to changes in human behaviour, demography, the environment, and/or the impact of (undocumented) interventions. However, such situations are obviously not stable in the long run.

Our study also shows that assortative mixing substantially influences the impact of interventions. Its importance may be even greater if mixing is correlated with MDA uptake, especially if high-risk groups are less likely to participate in MDA. If missed, such high-risk groups may reintroduce infection into the general population. As such, if assortative mixing occurs at a very local scale, e.g. at household level, high coverage of treatment within households may be even more important than overall population treatment coverage. Further, bounce-back of infection levels is relatively slower under assortative mixing than with homogeneous mixing and may therefore occur later than expected, a pattern similar to relatively slower outbreaks of malaria in populations where mixing is more assortative [15]. Therefore, identifying, treating, and monitoring of high-risk groups is highly important. Similarly, if vector control is considered, locating and targeting those breeding sites that are most important for transmission is pivotal. The same applies if low prevalences are sustained by movement of infected humans and/or flies over larger distances; uniform intervention coverage and in particular coverage of high risk groups/areas is pivotal to minimise the risk of recrudescence of infection after stopping interventions.

Unfortunately, proving existence and quantifying the degree of assortative mixing with data may not be easy. If assortative mixing plays a relevant role in helminth transmission, it is most likely related to patchy distribution of vectors or environmental reservoirs of infection. For example, onchocerciasis transmission in forest areas is sometimes driven by multiple smaller fly breeding sites. Because in savanna areas the number of fly breeding sites that a village is exposed to is typically limited, assortative mixing (if any) may be more likely to be driven by a sub-group of individuals (e.g. fishermen) that frequent a breeding site further away from the community. In both cases, local fly data from such areas may be informative. More specifically, locally high prevalence among flies and/or annual transmission potential (i.e. the number of fly bites times the average number of L3 larvae per fly bite) could perhaps be linked to a specific sub-group of humans that spend more time near certain fly breeding sites. In addition, data on the intensity distribution of infection in a community may provide some information in communities where prevalence of infection is under 30%, although subtle patterns may easily be masked by measurement and sampling error.

Eventually, genetic studies may provide an answer to the question who infects whom. Although such studies have not yet been attempted, genome-wide analyses of *Onchocerca volvulus* populations have been performed in Cameroon and Ghana, demonstrating that this technique is able to genetically distinguish geographically separate worm populations (i.e. populations that mix in a limited fashion) [28]. To what extent such analyses can be used to quantify the degree of past and ongoing mixing remains to be investigated. For soil-transmitted helminths and schistosomiasis, quantitative studies of human open defaecation may help inform the degree and importance of assortative mixing for transmission and impact. Although challenging to reliably quantify, questionnaires about or direct observations of where uniquely identified people defaecate exactly (preferably repeated over a period of time) could help quantify the spatial patchiness of transmission sites and how often they are frequented by whom, allowing construction of more realistic transmission models that account for assortative mixing.

We realise that our implementation of assortative mixing is a simplification of reality. In real-world situations more than two risk groups may well exist, and the degree of assortative mixing between such groups may differ from what we assume here. Still, a related modelling study on hepatitis C transmission in and between the general populations and high-risk groups demonstrated that simply adding the process of assortative mixing itself captures much of the qualitative behaviour of a system, and adding more risk groups to the system does not change its behaviour much [29].

In conclusion, assortative mixing could play an important role in helminth transmission dynamics, but is difficult to measure in real-world situations. The presence of assortative mixing will reduce the chance of achieving interruption of transmission. More detailed data on infection intensity distribution in human and vector populations (or environmental reservoirs), and actual contact rates between humans and vectors or environmental reservoirs are needed to answer to which extent assortative mixing plays a role in reality. For modelling studies, introducing the phenomenon of assortative mixing will help to explain low stable endemic situations.

---

**S1 Figure.**
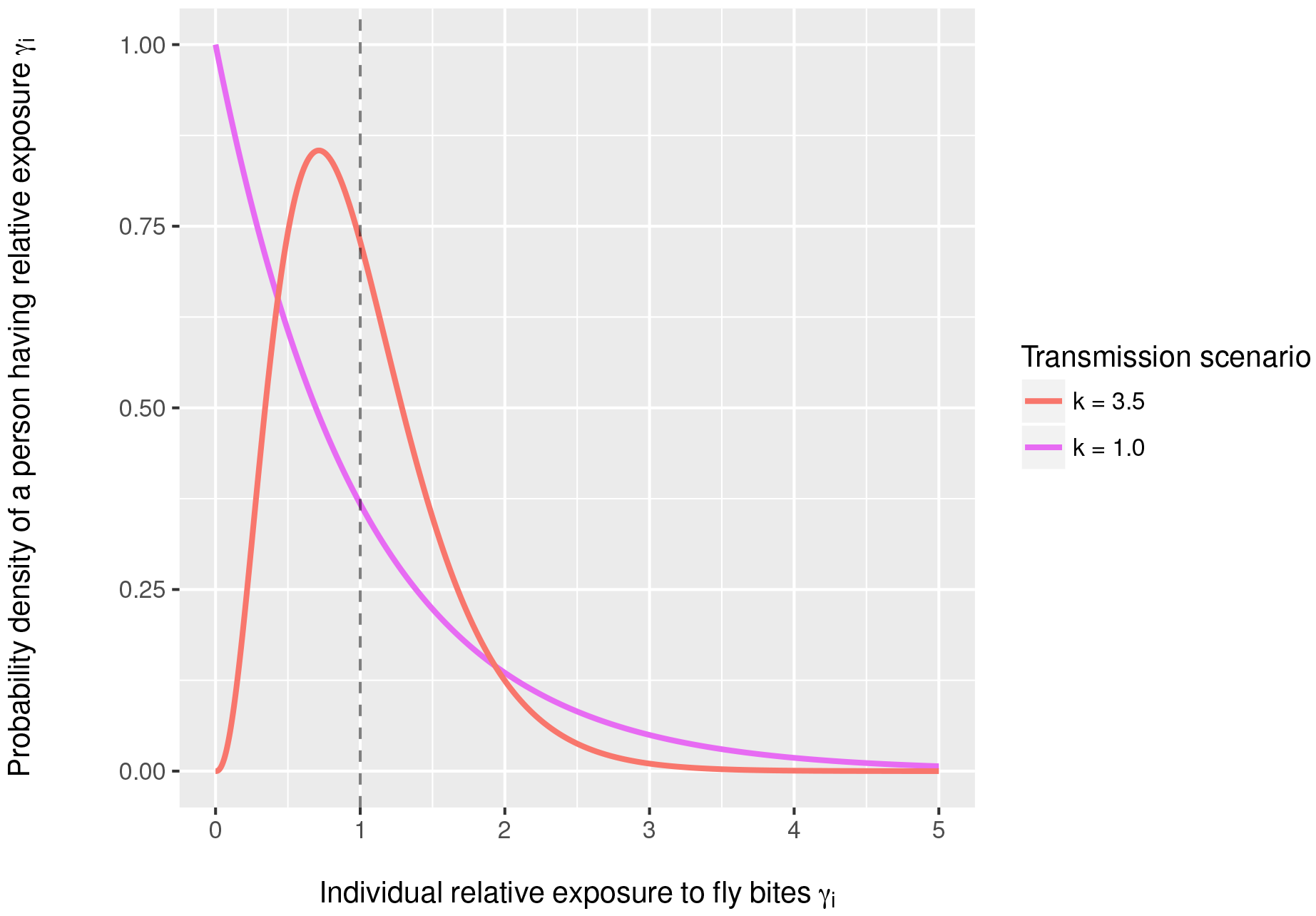
Assumed distribution of relative exposure to fly bites in the human population. Relative individual exposure to fly bites is assumed to follow a gamma distribution with shape and rate equal to *k* (3.5 or 1.0) and mean 1.0 (dashed vertical line).

**S2 Figure.**
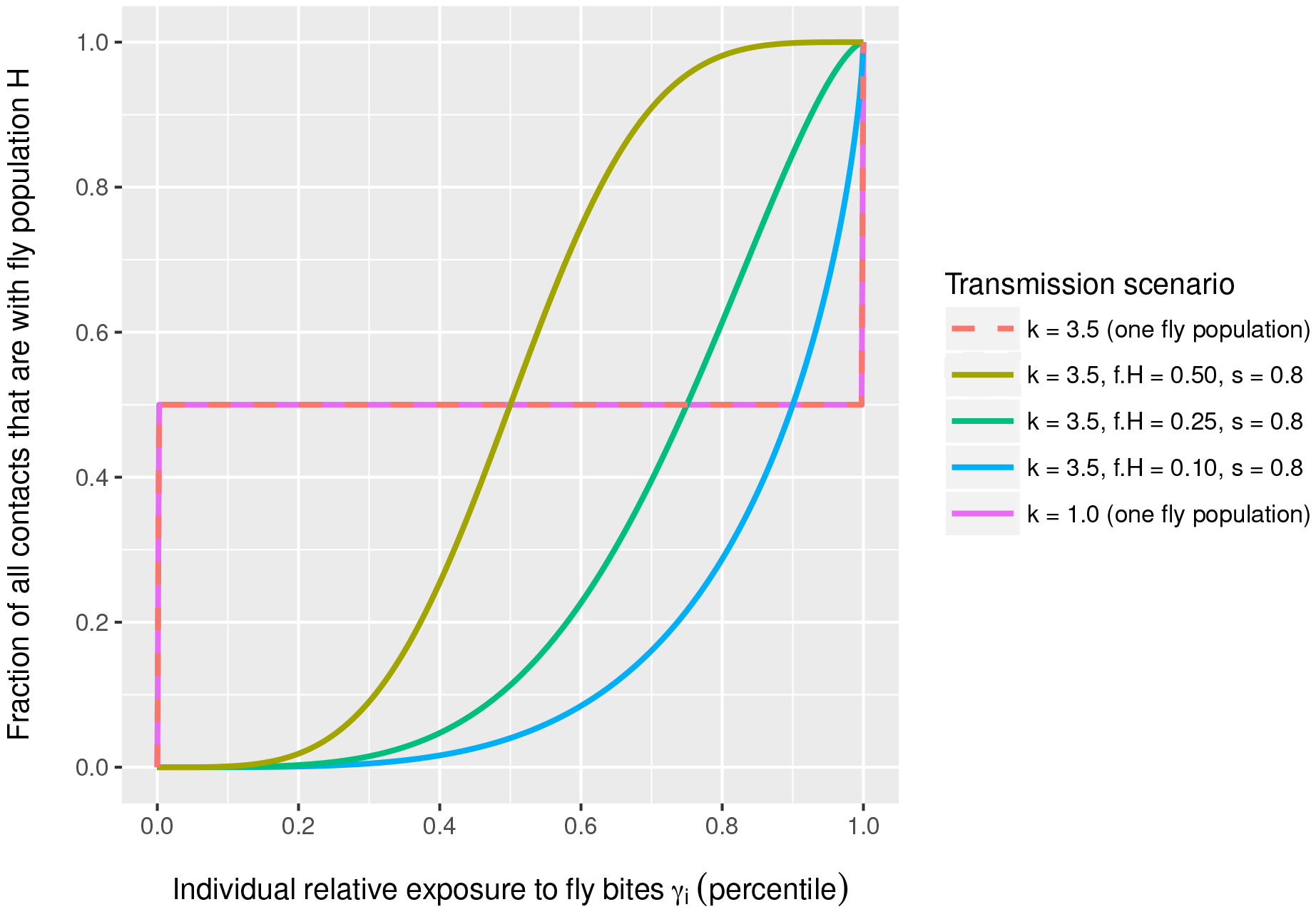
Assumed association between individual relative exposure to fly bites and fraction of bites received from fly population *H*. Red (dashed) and purple lines overlap perfectly because they both represent a setting where all individuals are equally exposed to the two fly populations in the model, which means that the two populations effectively function as a single fly population that mixes homogeneously with the human population.

**S3 Figure.**
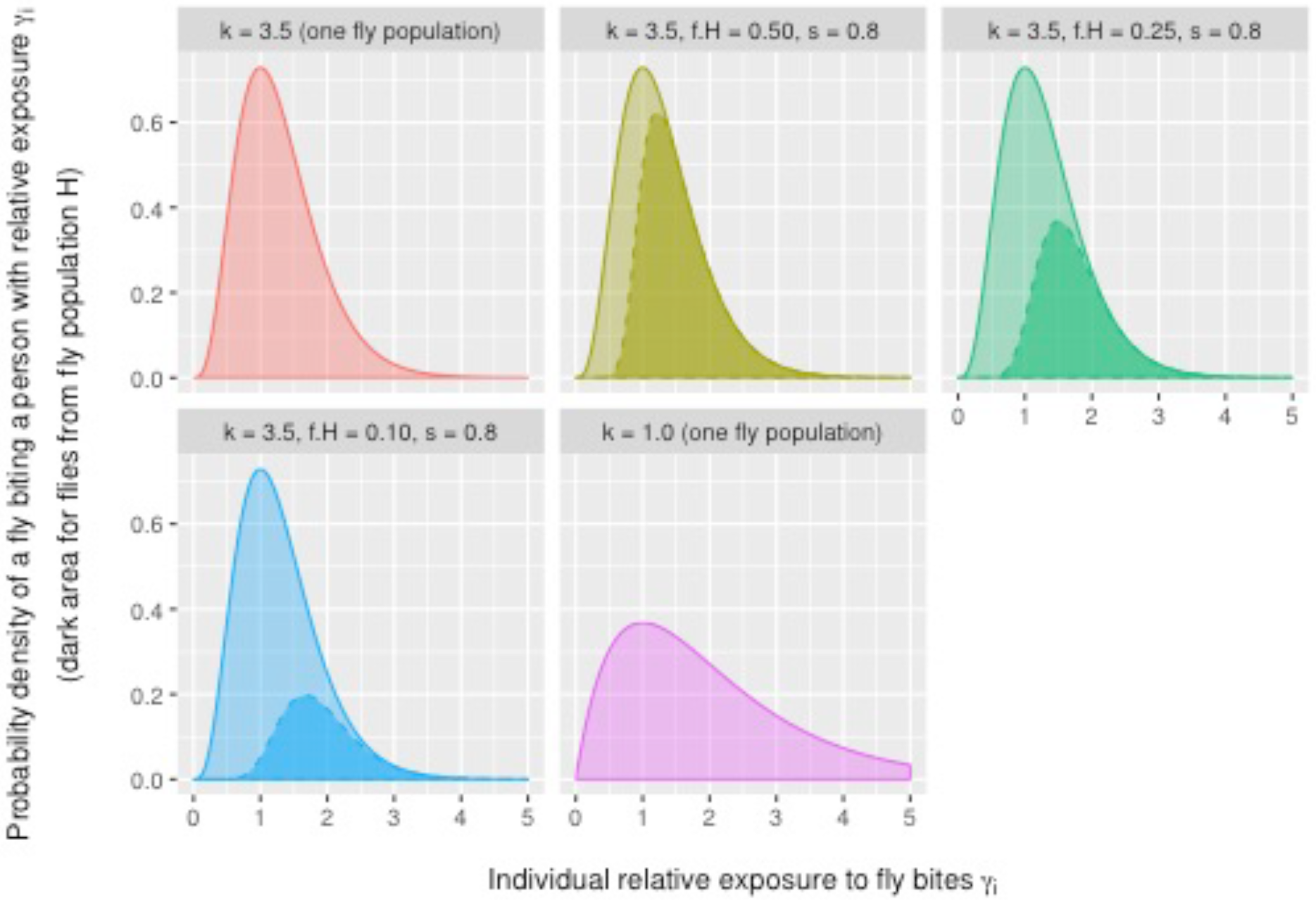
Probability density that a fly will bite a person with a given relative exposure for transmission scenarios with either one fly population (red and purple) or two fly populations (other colours). Darker areas represent bites by flies from fly population *H*.

**S4 Figure.**
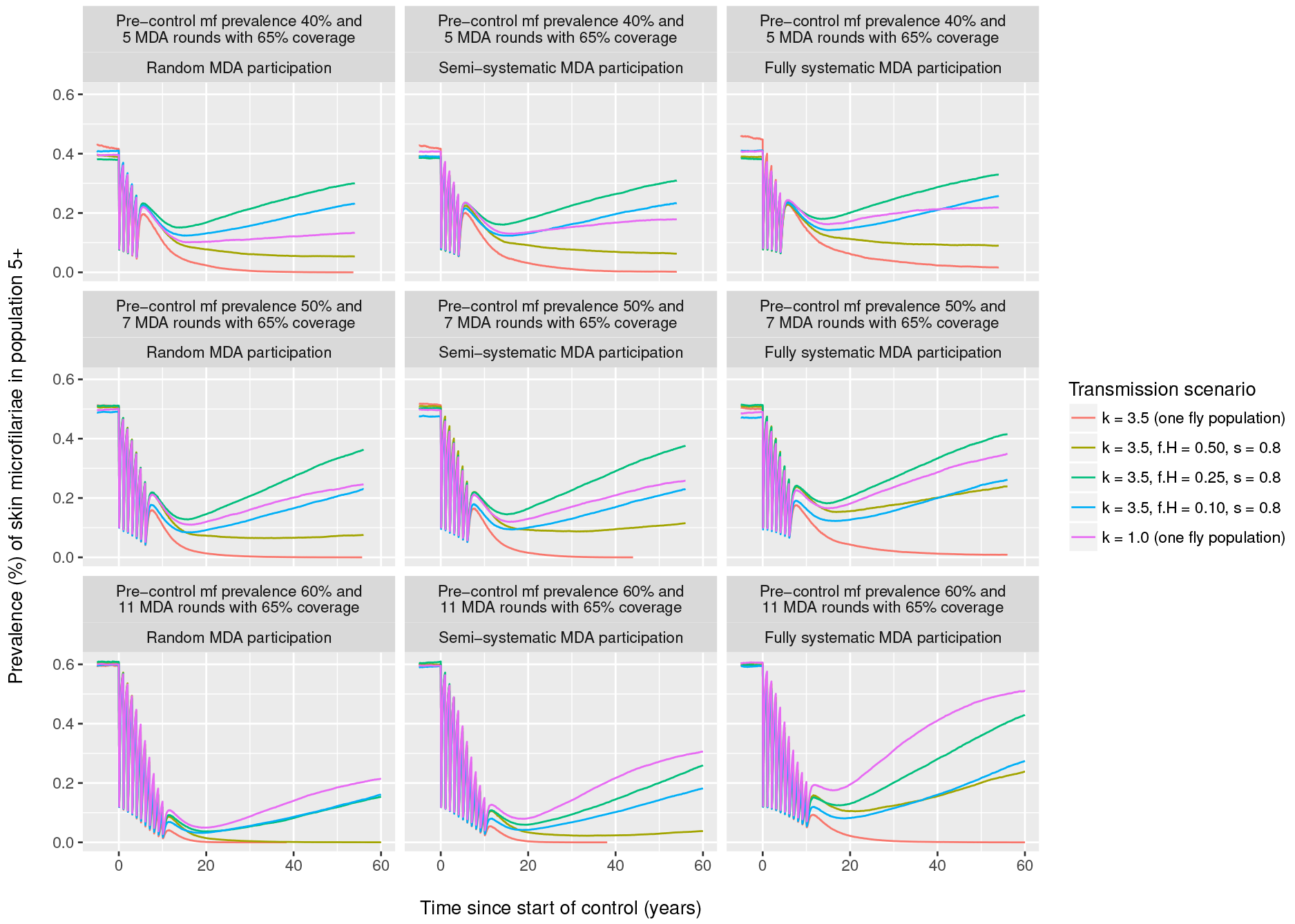
The influence of mixing patterns on trends in prevalence of skin microfilariae during mass drug administration. Simulations represent three setting with pre-control prevalence of about 40%, 50%, or 60% in the population of age 5 and above where 5, 7, or 11 rounds of mass drug administration rounds are implemented at 65% coverage of the general population (rows of panels). Columns of panels represent three different assumptions about patterns in MDA participation: completely random (participation to MDA is independent of past participation) vs. semi-systematic (as in Figure 2) vs. completely systematic (same individuals always participate). Lines represent the average of 200 repeated simulations, including simulations that resulted in elimination.

**S5 Figure.**
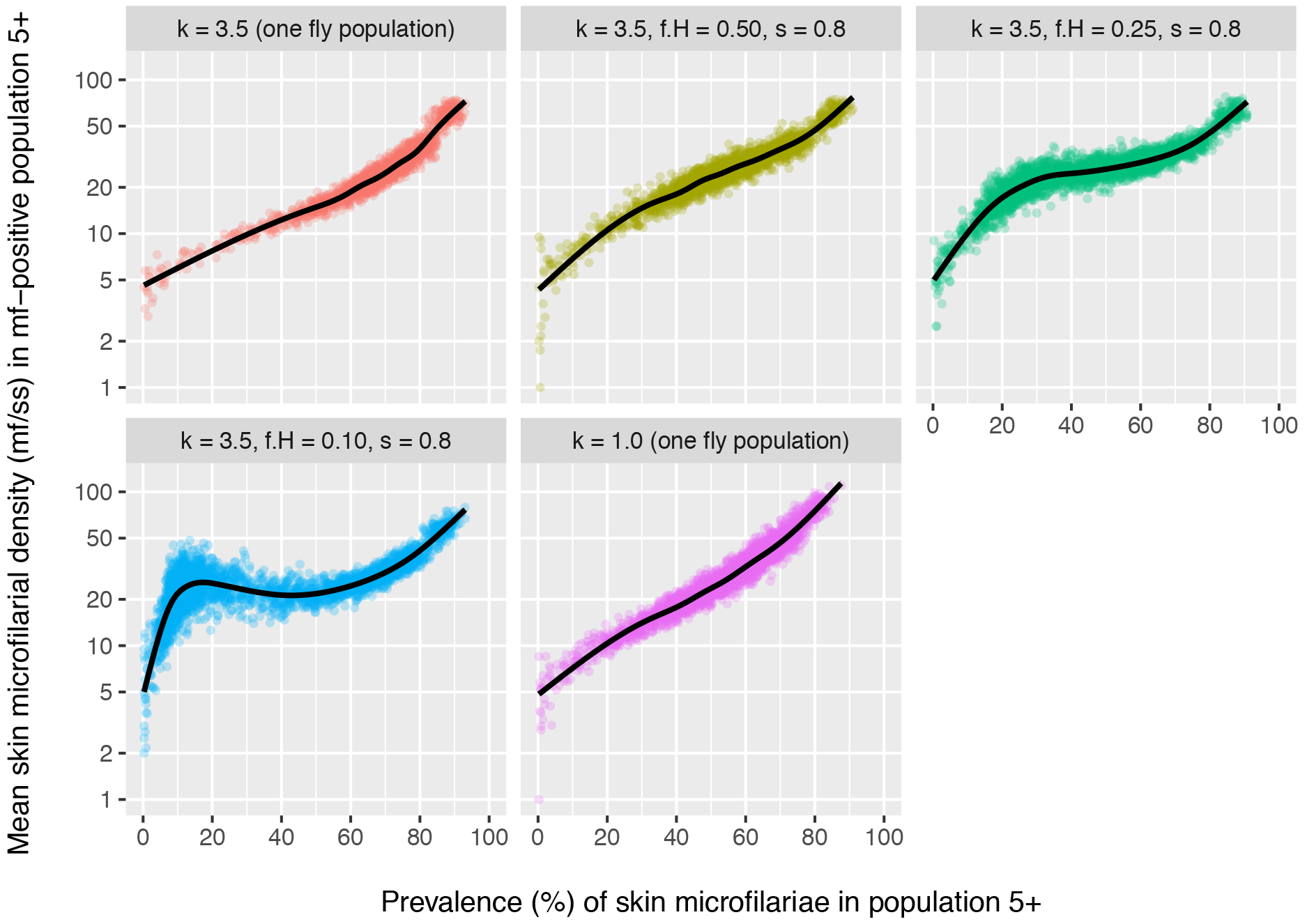
Pre-control density of mf in the skin among mf-positive individuals of age 5+ and above. Each bullets represents the result of a single simulation. Simulation were run using the same range and values of annual biting rate (ABR) as used in Figure 1, and for each value of ABR 150 repeated simulations were performed. Lines are based on a generalised additive model with integrated smoothness estimation, fitted to the individual bullets.

**S6 Model code.** R scripts to run the model and analyses.

